# Visual lateralisation in female guppies demonstrates social conformity but is reduced when observing a live predator

**DOI:** 10.1101/2025.05.06.652411

**Authors:** Iestyn L. Penry-Williams, Culum Brown, Christos C. Ioannou

## Abstract

Living in groups offers individuals a way of reducing their risk of predation. Visual lateralisation, characterised as an asymmetry in eye-use, may offer an additional advantage to group-living animals by enabling them to manage two concurrent visual tasks simultaneously. This could enhance multi-tasking efficiency by facilitating cohesion with group mates while monitoring for threats. In our study, we examined visual lateralisation of Trinidadian guppies (*Poecilia reticulata*) tested either alone or in groups, in either the presence or absence of a live predator, the blue acara (*Andinoacara pulcher*). We consistently observed low levels of visual lateralisation across all treatments. Contrary to our expectations, however, guppies exhibited significantly higher absolute lateralisation when tested alone in the absence of the predator compared to the other treatments. Moreover, a significant left-eye bias was observed when the predator was present, and the fish showed a right-eye bias when the predator was absent. Use of a repeated measures design and assessing individual and group ID as random effects provided evidence that both relative and absolute laterality were repeatable at the group level, but there was limited evidence for repeatability at the level of the individuals. This repeatability in lateralisation when tested as a group, but not when individual fish composing these groups were tested alone, suggests social conformity in lateralisation. Our results suggest that social processes may have a significant impact on within-population variation in lateralisation.

## Introduction

Predation is a major driver of group formation in prey species. Individuals in groups often experience higher survival compared to solitary individuals through mechanisms including risk dilution, the avoidance effect, the confusion effect and group vigilance (Ioannou 2021)□. Maintaining visual contact with other group members to sustain group cohesion while staying vigilant to threats is required to maximise an individual’s chance of survival (Ward et al., 2011)□. However, performing two concurrent visual tasks may hinder performance of both tasks unless visual information can be efficiently processed (Dadda & Bisazza, 2006a; Dukas, 2004; Rogers et al., 2013)□.

Shoaling in fish is notably synchronous, particularly in environments with higher predation, where shoaling is more prevalent and social tendencies of individuals (i.e. “sociability”) are heightened (Gartland et al., 2022; Kelley & Magurran, 2003; Seghers, 1981)□. Individuals in groups might derive further advantages by exhibiting a directional bias (left or right asymmetry) in certain behaviours, known as lateralisation or “handedness” (Rogers et al., 2013). For example, some visual tasks may be preferentially performed with a certain eye (i.e. “visual lateralisation”), or an individual may demonstrate a consistent directional response (i.e. “motor lateralisation”). This is believed to be linked to a cerebral partitioning of cognitive functioning (Hulthén et al., 2021)□. Behavioural lateralisation is typically assessed through “relative laterality”, i.e. the directional bias of the individual (left or right), and “absolute laterality”, the strength or intensity of this bias regardless of directionality. Behavioural lateralisation has been observed across a range of taxa and behaviours including tool use, predator avoidance, and escape responses (Rogers et al., 2013)□. Lateralised individuals have demonstrated greater escape responses (Dadda et al., 2010)□, greater foraging capabilities (Dadda & Bisazza, 2006b)□, and tend to occupy safer positions in a group when under predator presence (Bibost & Brown, 2013; Middlemiss et al., 2018)□. However, despite the potential advantages of lateralisation, it has not been consistently observed in prior studies, with variability in both the directionality and intensity within and between populations (Bisazza, Rogers, et al., 1998; Penry-Williams et al., 2022; Roche et al., 2020)□.

In environments with high levels of predation, individuals are expected to demonstrate enhanced visual lateralisation, an asymmetric bias in eye-use when viewing a stimulus, which can be assessed through both relative and absolute lateralisation. Fish species at lower trophic levels typically have laterally positioned eyes, offering limited binocular overlap (Vanegas and Ito, 1983). This configuration means that stimuli are predominantly viewed by only one eye at a time (Middlemiss et al., 2018; Vanegas & Ito, 1983)□. The information gathered from each eye is primarily processed by the contralateral hemisphere, allowing for the potential division of two concurrent visual tasks between brain hemispheres if one eye is used for each task (Bisazza & Brown, 2011; Dadda et al., 2009)□. Cognitive partitioning, such as this, would enable more efficient information processing and multi-tasking (Bisazza & Brown, 2011; Miletto Petrazzini et al., 2020; Vanegas & Ito, 1983)□. This is particularly beneficial in the context of shoaling, as an individual could monitor shoal mates while simultaneously surveying other external stimuli, such as predators or food (Bisazza & Dadda, 2005). If specialisation of a visual task was dedicated to a particular eye, then detection latency and neural processing time would be minimised, allowing for an increased response efficiency (Bisazza et al., 2000; Brown et al., 2004; Dadda & Bisazza, 2006a; Vallortigara, Rogers, et al., 1999)□. Furthermore, if multiple individuals within a shoal were to demonstrate lateralisation, this could enhance group synchronisation, cohesion, and escape capacity (Brown, 2005; Brown et al., 2004; Frasnelli & Vallortigara, 2018; Miletto Petrazzini et al., 2020)□. In such circumstances, a mixture of individuals displaying right- and left-alignment may be expected dependent on their shoal positioning, as has been found in crimson-spotted rainbowfish (*Melanotaenia duboulayi*) and black-banded rainbowfish (*Melanotaenia nigrans*) (Bibost & Brown, 2013).

Lateralised eye-use in fish has been extensively documented across various species and contexts, particularly in response to predatory and social stimuli. For instance, studies have shown that *Brachyraphis episcopi* (Brown et al., 2004), goldbelly topminnows (*Girardinus falcatus*) (Facchin et al., 1999),LJ Bahamas mosquitofish (*Gambusia hubbsi*) (Hulthén et al., 2021), and eastern mosquitofish (*Gambusia holbrooki*) (Bisazza et al., 1997a) all exhibit lateralised eye use when exposed to predatory stimuli □. In Trinidadian guppies (*Poecilia reticulata*), even exposure to a predator’s odour alone was sufficient to induce lateralisation (Broder & Angeloni, 2014)□. This phenomenon extends to social interactions as well, with asymmetrical eye-use observed in poecilids (Fuss et al., 2019; Sovrano et al., 1999) and several other fish species (Sovrano et al., 2001) when viewing conspecifics or their own reflections. □

However, most of these studies have focussed on assessing lateralisation in solitary individuals (Broder & Angeloni, 2014; Facchin et al., 1999; Sovrano et al., 1999)□, despite many species, including guppies, often occurring in social groups. Furthermore, while some research has explored population-level lateralisation in dichotomous “high versus low” predation environments (Brown et al., 2004), few studies have directly examined how the presence of a live predator influences lateralisation in real-time, particularly in a social context. The gap in our understanding is significant, as the benefits of lateralisation are theorised to be especially pronounced in group-living species facing predation pressure (Brown et al., 2004).

Recent work has begun to address some of these limitations. Johnson et al. (2020) working with *Xenophallus umbratils* included a predator as a stimulus in their arena trials and observed a strong impact on lateralisation. However, *Brachyrhaphis rhabdophora* have demonstrated no significant difference in laterality based on predation regime, even when faced with a predatory stimulus (Callaway et al., 2023). These contrasting findings highlight the variability in lateralisation responses across species and contexts, emphasising the need for more comprehensive studies that consider multiple factors simultaneously. To maximise predator evasion and escape, visual lateralisation is believed to be most efficiently adopted in conjunction with shoaling behaviours (Bibost and Brown, 2013; Bisazza et al., 2000). If lateralisation is beneficial as an anti-predator behavioural mechanism, then we should expect this to be consistently demonstrated when under predatory threat due to the severe fitness consequences of predation (Toscano et al., 2014)□. While evolutionary history may select for lateralisation, plasticity in displaying lateralisation is influenced by individual experience and environmental context. Determining the conditions under which lateralisation is expressed and whether lateralisation is consistent and repeatable is fundamental to determining the ecological drivers of lateralisation and in predicting how prey species may respond to potential threats (Killen et al., 2016)□. Our study aims to address these limitations by investigating visual lateralisation in guppies under more ecologically relevant conditions. We used the predator-prey model system of female Trinidadian guppies (*P. reticulata*) and the predatory blue acara (*Andinoacara pulcher*) to investigate the interaction between predation risk, shoaling, and visual lateralisation in prey. We assess both relative and absolute lateralisation of each guppy in multiple treatments. By examining lateralisation both in solitary fish and in groups, and in the presence and absence of a live predator, we can better understand how social context and predation threat interact to influence lateralisation. This approach allows us to test whether the proposed benefits of lateralisation for group living and predator avoidance are realised under more ecologically realistic conditions.

The first aim of this study was to validate that the guppies perceived the predatory stimulus as a threat. This was assessed by measuring the guppies’ “attack cone avoidance” around the acara; i.e. the guppies should avoid being directly in front of the acara’s head if they perceive this stimulus as a potential threat (Magurran & Seghers, 1990)□. Next, the second aim was to determine whether shoaling and/or a live predatory stimulus impacts visual lateralisation. We hypothesised that both maintaining shoaling behaviours in a group and viewing a live predator should enhance lateralisation, with a potential additive effect when multi-tasking both activities. The final aim was to assess the repeatability of visual lateralisation (Penry-Williams et al., 2022; Vinogradov et al., 2021)□. We hypothesised that if visual lateralisation is an adaptive anti-predatory trait, then this behaviour should be more consistent and repeatable when fish are exposed to the predatory stimulus than when they are not (Toscano et al., 2014)□.

## Materials and methods

### Study species: Trinidadian guppies (*P. reticulata*)

Trinidadian guppies have been previously used as a model organism for investigating the effect of predation risk on lateralisation (Broder & Angeloni, 2014; De Santi et al., 2000; Irving & Brown, 2013)□. The population used in this investigation were descendants of wild guppies collected from a high predation environment in the Guanapo river in Trinidad (Moonan: 10.6082°N 61.2547°W) in April 2019 by the Guppy Project (University of Oxford). Guppies were exported to the John Krebs Field Station (Oxford, UK) where they were selectively bred across three generations to prevent inbreeding and maintain genetic diversity. Guppies were maintained between 25 to 27°C and fed twice daily with either live brine shrimp nauplii or liver paste, with a 12:12 light:dark photoperiod. Guppies were transferred to the University of Bristol (Bristol, UK) in December 2020 by car for approximately 1.5 hours. All fish were alive and in good condition upon arrival to the University of Bristol.

Preceding investigation, guppies were maintained in mixed-sex groupings of approximately 50 – 100 individuals in 90 L holding tanks (length x width x height: 70 x 40 x 35 cm) furnished with plastic foliage and a sand substrate. Guppies were fed once per day ad libitum with either brine shrimp or fish flake and maintained at 26 to 28°C and a 12:12 light:dark photoperiod. Guppies used in this investigation had previously been tested in a study exploring the short-term effects of temperature and turbidity on social behaviours in February and March 2021 (detailed in Allibhai et al., 2023), at least two months prior to use in the current study.

Only female guppies were used for this experiment to standardise the experimental protocol due to their stable social interactions (Croft, Morrell, et al., 2006)□, and to exclude sexual behaviours and harassment from males (Cummings, 2018)□. Female guppies were sorted into size-selected groups (n_grp_ = 32) to recognise individuals across trials without the need for elastomer tagging.

Guppies were measured (mean standard length ± SD: 26.1 ± 5.7 mm) and sorted into size classes: small (< 22 mm), medium (23 – 29 mm) and large (> 30 mm). One individual from each size class was haphazardly selected to form each group of three fish (mean group length ± SD: 26 ± 1 mm). Small deviations from size classifications were made in a few instances where fish in a particular size class had limited availability (n_ind_ = 6). Groups were viewed on the camcorder used for the trials to ensure individuals could be recognised and identified within their groups. Groups were housed in separate breeding nets (length x width x height: 16 x 13 x 13 cm) for the duration of the experimental period (18 days) to maintain group identities (n_grp_ = 32) with two breeding nets per 45 L holding tank (length x width x height: 35 x 40 x 35 cm). Breeding nets were kept in tanks on a filter system to ensure adequate water quality, oxygenation, and to prevent temperature fluctuations. Groups were acclimatised for 72 hours prior to testing. In instances where an individual within a group died during the experiment, they were replaced with a size-matched individual (n_ind_ = 12). The new individual and group were assigned a new individual ID and group ID, respectively, and acclimatised for 24 hours prior to testing.

During the investigation, all water quality parameters (pH, ammonia, nitrates and nitrites) were within the recommended range for the species and were monitored on the recirculated filter system weekly. Guppies were fed commercial fish pellets at 17:00, approximately 16 to 24 hours before trials.

### Study species: blue acaras (*A. pulcher*)

Blue acaras are a major predator species of Trinidadian guppies (Deacon et al., 2018; Zanghi et al., 2024)□. A total of fourteen blue acaras were used as the predatory stimulus to induce a heightened perception of predation risk (mean standard length ± SD: 86 ± 11 mm). Acaras were held in a 90 L holding tank (length x width x height: 70 x 40 x 35 cm) furnished with plastic foliage, plastic tubes, and a sand substrate base. Acaras were fed commercial fish pellet at 17:00, approximately 16 to 24 hours before trials. Acaras were maintained at a temperature of 26 to 28°C and a 12:12 light:dark photoperiod. Individual acaras were used in a maximum of one trial each day to limit potential stress.

### Lateralisation assays

Trials took place in two testing blocks in June and July 2021. Treatments manipulated both whether the predator was present or absent, and whether guppies were tested alone (n_ind_ = 96) or in their group (n_grp_ = 32) in a fully-factorial design; the treatments were “Predator-Solitary”, “Predator-Group”, “Control-Solitary” and “Control-Group”. Each individual was subject to all four treatments twice (i.e. 8 trials total), with the treatment order following a Latin-square design. Two trials were run in parallel, with a “Predator” trial and a “Control” trial taking place concurrently. “Group” and “Solitary” treatments took place on alternating days. Control and Predator treatments were alternated between the two experimental set-ups after assaying each group to account for potential asymmetries; whether the trial took place in the left or right set-up was included in the statistical analyses. The testing order of fish groups was randomised each day using random.org. In the Solitary treatment, the testing order of individuals within each group was also randomised. Solitary trials occurred between 9:00 to 17:00 (n = 24 trials per day). To accommodate for the fewer trials on days that Group trials took place (n = 8 trials per day), start times were varied. Group trial start times were randomly determined each day to fall within the windows of 9:00 to 11:00, 11:00 to 13:00, or 13:00 to 15:00. Fish were used in a maximum of one trial each day to limit potential stress resulting from the experimental procedure.

The apparatus used in this experiment was similar to that used by Broder & Angeloni (2014) and Zanghi et al. (2023)□. The assay consisted of a white circular container (*Figure 1*: diameter x height: 32 x 30 cm) with a central transparent chamber containing either a blue acara (Predator treatment) or left empty (Control treatment). The predators were restricted to the central compartment to avoid olfactory cues from the predator and prevent direct physical contact between predator and prey. Groups, or a solitary individual, were introduced into an opaque white acclimatisation section (white PVC tube: diameter x height: 5.5 x 10 cm) in the area surrounding the central chamber for ten minutes preceding the trial, after which the video camera was started (Panasonic HC-X920: 1920 x 1080 pixel resolution, 50 frames/second), and the acclimatisation tube was lifted. Trials lasted for 15 minutes. All tanks were shaded from direct light using a translucent plastic sheet to prevent a light-induced turning bias and to reduce reflections on the surface of the water to facilitate computer tracking from video. Water temperature was between 26 to 28°C and was replaced from the filtration system following each trial. Water level was maintained at 8 cm in the outer ring, and 11 cm within the central chamber.

**Figure 1:**
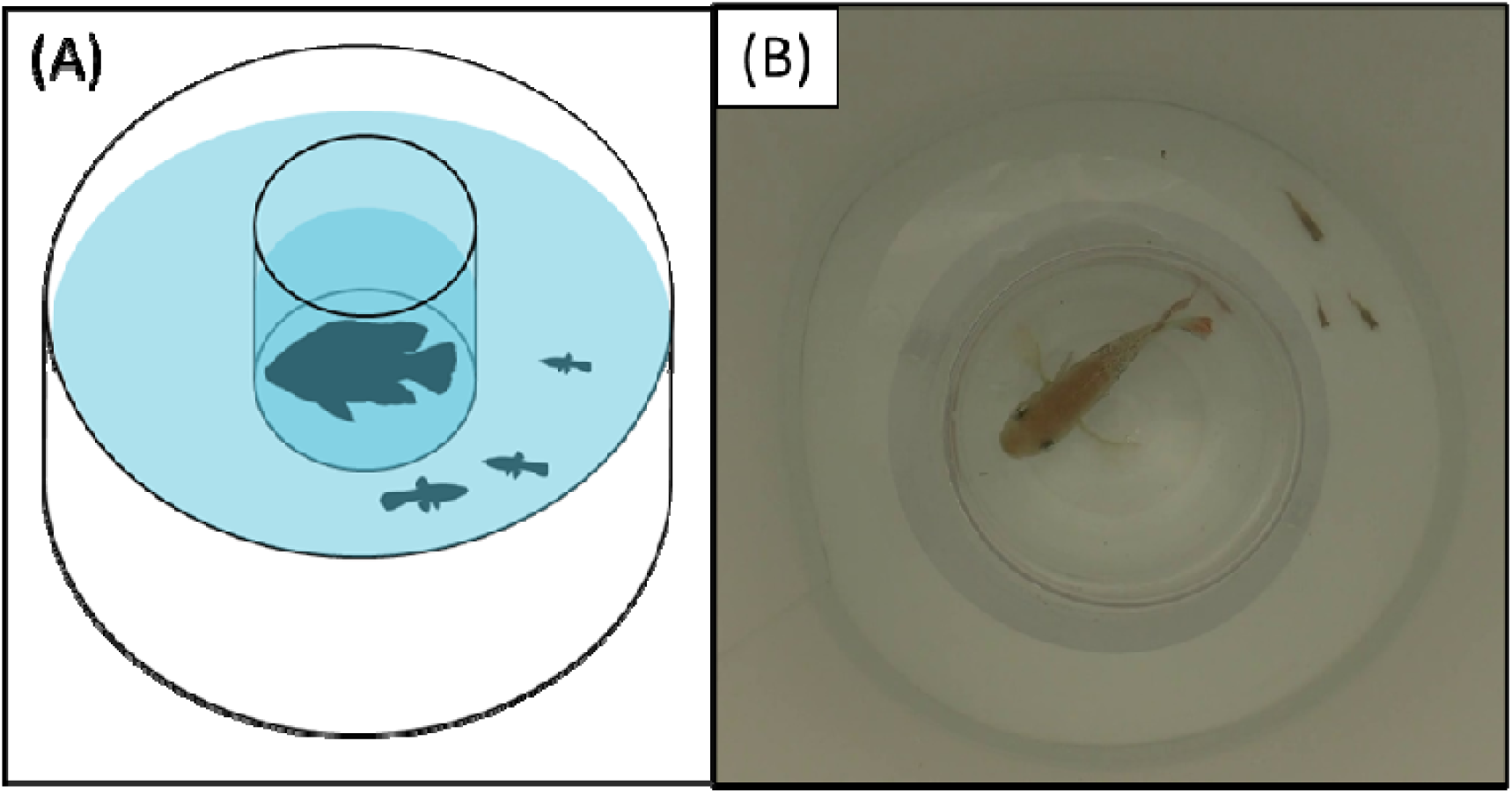
**(A)** Schematic representation of the lateralisation assay displaying a group trial with the predator stimulus within the chamber. **(B)** Still image from a video of a trial demonstrating the lateralisation assay, blue acara predator (*A. pulcher*), and displaying size differences between the individual guppies (*P. reticulata*) within the group.

### Data processing: guppy trajectories

All trial videos were converted from .MTS format to .mp4 (1920 x 1080 pixel resolution, 30 frames/second) using Handbrake (v1.2.0) (The Handbrake Team, 2018)L to facilitate tracking in idTracker (v2.1) (Pérez-Escudero et al., 2014)L. Following tracking, trajectories were manually assessed for accurate identification of guppies and their trajectories. Where inconsistencies were found, tracking parameters were adjusted, and tracking was reperformed. The centre and diameter of the central chamber was manually measured in ImageJ (v1.52a) (Rasband, 2018) from still images of each trial obtained using VLC media player (v3.0.8 Vetinari) (VideoLan, 2006)L. Data cleaning involved the removal of datapoints exceeding maximum speeds for Trinidadian guppies (140 cm/sec burst speed) to account for potential tracking errors (Chappell & Odell, 2004; Oufiero & Garland, 2009)L. Data were then interpolated using the na.interpolation function (imputeTS package) (Moritz & Bartz-Beielstein, 2017)L using a linear option and allowing a maximum gap of ten successional missing points (approximately 0.33 seconds) in R (v3.6.1) (R Core Team, 2019)□ with RStudio (v1.2.1335) (RStudio Team, 2019)□.

For each frame of video (30 frames/sec for 15-minute videos), relative lateralisation was calculated using χ = arcsin(sin(θ−□)), in which θ was the angle of the fish between two frames and □ is the angle of the arena radius through the position of the fish (Herbert-Read et al., 2015; Penry-Williams et al., 2022)□. Lateralisation indexes are relative to the centre of the central chamber. A score of χ > 0 demonstrates a clockwise orientation, i.e. using the right eye to view the centre of the arena, while a score of χ < 0 is an anti-clockwise orientation, i.e. using the left eye to view the centre of the arena. While the number of frames demonstrating right and left eye-use were used in the statistical analysis to assess relative lateralisation, to display the data these were used to calculate a relative laterality (RL) index (on a scale of -1 to 1) using RL = ((N_right_ − N_left_) / (N_right_ + N_left_)), where N_right_ is the number of frames of right-eye use and N_left_ is the number of frames of left-eye use. To calculate absolute laterality (AL) to assess the intensity of lateralisation, the absolute values of the relative laterality index were calculated, so that absolute laterality ranged from 0 to 1. Each individuals’ activity was calculated as their speed (cm/min) from the total distance travelled by the fish (cm) and dividing by the trial duration (min).

### Data processing: acara orientation and attack cone avoidance

An assumption of our study is that the guppies perceive the blue acara as a predatory threat. Therefore, in trials with a predator, whether the guppies avoided the attack cone of the acara was assessed, i.e. whether the guppies avoided being directly in front of the acara’s head (Magurran & Seghers, 1990)□. This was measured by assessing the guppies’ position in relation to the acara’s body orientation.

An automated custom ImageJ plugin using a convolution-based approach was used to detect the centre of mass and orientation of the acara (details available in Heathcote et al., 2020)□. The extracted angle and centre of mass was used to calculate the relative positioning of each guppy for each extracted frame of the trial, with 0° being directly in front of the acara’s head and 180° behind the acara’s tail. The activity of the acara was also assessed through the change in degrees per minute.

To facilitate tracking, the resolution and frame rate of the trial videos were reduced using ffmpeg (640 x 360 pixel resolution, 3 frames/second) (FFmpeg Developers, 2021)□. This reduction for evaluating the orientation of the acara did not impact the results. To verify the accuracy of the tracking, a subset of the trial videos (n = 10) was randomly selected and run at the full resolution (1920 x 1080 pixels resolution) and full frame rate (30 frames/second). The angles extracted from the full resolution and frame rate videos were found to be in close agreement with those from their downsampled version. The mean ± SD absolute difference in degrees was 0.10° ± 0.31° for resolution and 0.70° ± 0.67° for frame rate. Further comparisons of the guppies’ positioning relative to the acara’s head also showed good agreement between the full and downsampled videos. The relative difference between the output datasets, calculated as (full frame rate / reduced frame rate) x 100, had a mean ± SD of 99.75 ± 0.49% for resolution and 100.03 ± 4.06% for frame rate, indicating a high level of consistency between the full and downsampled datasets.

## Data analysis

All statistical analysis was performed in R (v3.6.1) (R Core Team, 2019)□ with RStudio (v1.2.1335) (RStudio Team, 2019)□. To validate guppies perceiving the blue acara as a predatory threat, it was initially assessed whether there was attack cone avoidance around the head of the acara, in trials with a predator present (Magurran & Seghers, 1990)□. Individual guppy positioning relative to the acara (median degrees from the acara’s head) was analysed as a response variable using a linear mixed-effects model (LMM). Explanatory variables were grouping treatment (“solitary” or “group”), guppy size (mm), guppy activity (cm/min), replicate number (first or second time completing that treatment), time of day, date, trial number (one to eight), side of assay (left or right set-up), testing block (first or second set of groups), predator activity (degrees turned/minute), and predator size (mm), with individual ID nested within group ID as the random effect.

Visual lateralisation was then investigated across all trials. Absolute laterality indexes (AL) were rescaled from 0 – 1 to 0 – 100 and rounded to the nearest whole number to fulfil the assumptions for a negative binomial generalised linear mixed-effects model (GLMM) using the glmer.nb function (lme4 package) (Bates et al., 2015)□. Relative lateralisation was assessed by using the proportion of frames with left versus right eye-use in a binomial GLMM using the glmer function (lme4 package). To fulfil model assumptions, the proportion of right versus left eye-use over each trial (27,000 frames) was scaled to proportions of 100. Grouping treatment (“solitary” or “group”), predation treatment (“predator” or “control”), guppy size (mm), guppy activity (cm/min), replicate number (first or second), time of day, date, trial number (one to eight), side of assay (left or right set-up), and testing block (first or second) were included as main effects, with individual ID nested within group ID as the random effect. For all models, the main effect with the highest p value (when > 0.05) was removed from each iteration of the model (i.e. backwards selection) and the model was re-run until only significant main effects remained. Grouping treatment and predation treatment were initially included as an interaction term, however, this interaction was removed if this had the highest p value (> 0.05), which was the case in all models. Grouping treatment and predator treatment were not removed from the models as explanatory terms at any stage as these are integral to the hypotheses being tested.

Data from trials with a predator were also analysed separately to assess whether additional factors significantly impacted guppy behaviour. In these models, reflecting those previously described, predator activity (degrees turned/minute), guppy positioning relative to the acara (median degrees from the orientation of the acara’s head), and predator size (mm) were added to the models as main effects along with the aforementioned main and random effects, and excluding the predator treatment explanatory variable. The assumptions of all models were verified with QQ plots and residuals versus fitted values using the residual diagnostics for hierarchical (multi-level/ mixed) regression models (DHARMa) package (Hartig, 2019)□. Variance inflation (multicollinearity) was assessed using the vif function (car package) (Fox & Weisberg, 2011)□. Individuals not completing a minimum of one of each of the treatments were excluded from the analysis (N_ind_ = 12, N_obs_ = 15).

Repeatability (R) in guppies’ behavioural traits (positioning relative to the acaras’ head, absolute laterality, and relative laterality) were assessed using repeatability estimates with parametric bootstrapping at 1,000 iterations (rptR package) (Stoffel et al., 2017)□. This was initially run on all trials together, and then on subdivisions of each treatment (“Predator-Solitary”, “Predator-Group”, “Control-Solitary”, and “Control-Group”). Median degrees from the acara’s head was assessed with a Gaussian datatype option. For absolute lateralisation, individuals were ranked (with averaged tied ranks) to avoid the violation of homogeneity of variances in the model’s residuals and an estimate of repeatability assessed via a Gaussian datatype option. Relative lateralisation was assessed through the proportion of frames with left versus right eye-use (left and right) using the proportion datatype option. For all models, repeatability in both individual ID and group ID was investigated. LMMs and GLMMs with the same model structure were fitted to check and verify the model assumptions.

### Ethical note

All experimental procedures and housing conditions were approved by the University of Bristol Animal Welfare and Ethical Review Body (UIN/21/003). All fish were monitored during the experimental period to ensure that they did not display overt signs of stress, and after testing were retained in the laboratory for use in future experiments.

## Results

### Attack cone avoidance

The guppies were consistently found to display attack cone avoidance (measured by median degree from the acara’s head), positioning themselves at 123° ± 10° (mean ± SD) for the solitary trials and 125° ± 7° for the group trials (*Figure 2*). Guppies spent 73.18 ± 7.50% (mean ± SD) and 71.42 ± 5.60% of the trial behind the predator (> 90°) for the solitary and group trials, respectively. This evidence indicates that guppies perceived the acara as a predatory threat in both solitary and group trials (Magurran & Seghers, 1990)L. No explanatory variable significantly affected the guppies’ attack cone avoidance, including whether the guppies were tested alone or in groups (*Figure 2*: Grouping treatment: LMM: χ^2^ = 1.94, p = 0.163).

**Figure 2:**
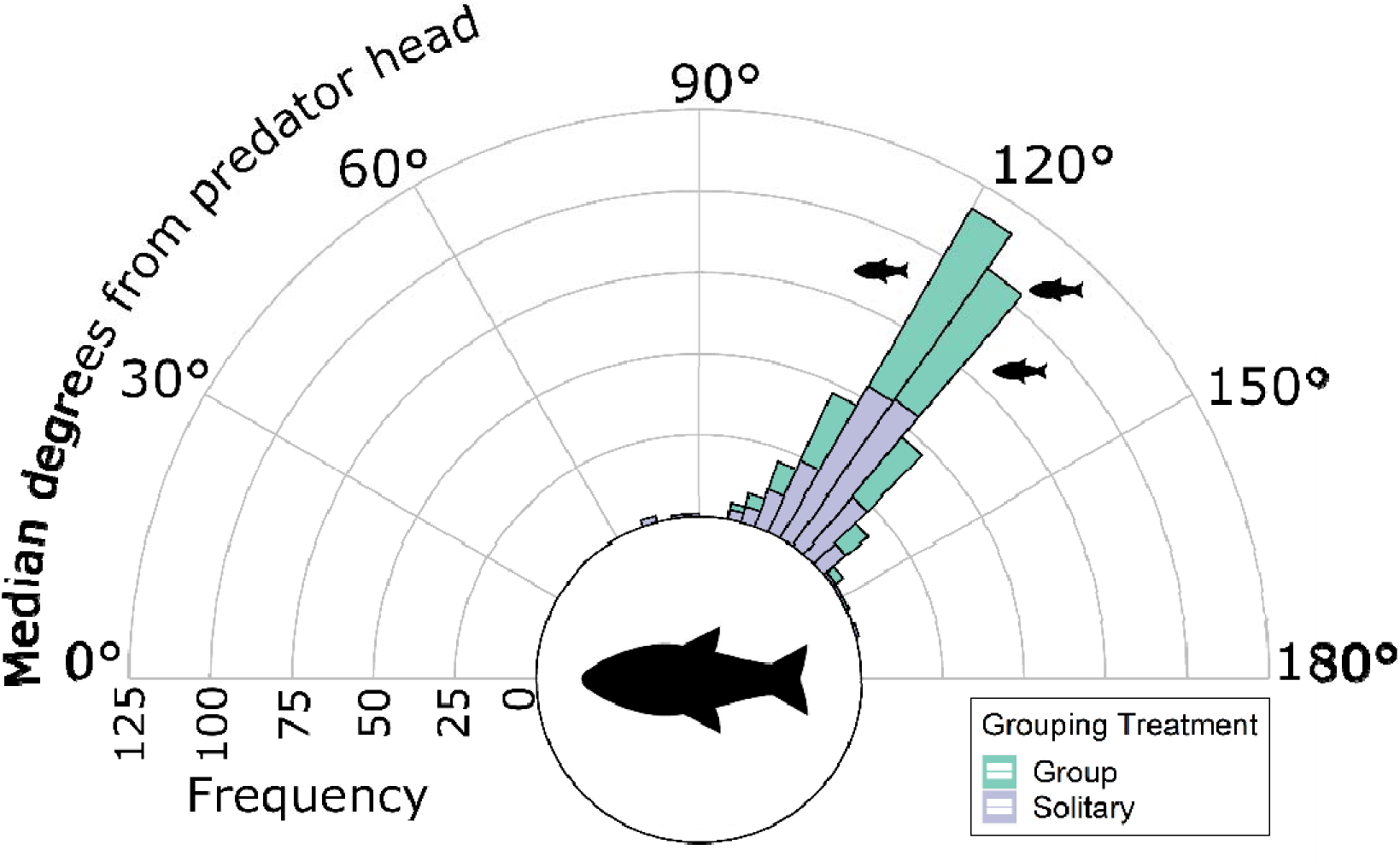
Stacked histogram of the median degrees of the guppies’ position from the acara’s head, with data split by the shoaling treatment (Blue = Group, Purple = Solitary). The large fish icon indicates orientation of the blue acara (*A. pulcher*), smaller fish icons represent the Trinidadian guppies (*P. reticulata*).

Across treatments, repeatability in attack cone avoidance (measured by median degrees from the acara’s head) was not found within individuals (LMM: R = 0.023, 95% CI = 0 – 0.113, N_obs_ = 379, N_ind_ = 97, p = 0.301), nor within their respective groups (R = 0.034, 95% CI = 0 – 0.29, N_obs_ = 379, N_grp_ = 38, p = 0.132). However, Group ID did produce significantly repeatable attack cone avoidance behaviour within the “Predator-Group” treatment (*Table 1*), as well as individual ID for the “Predator-

**Table 1:**
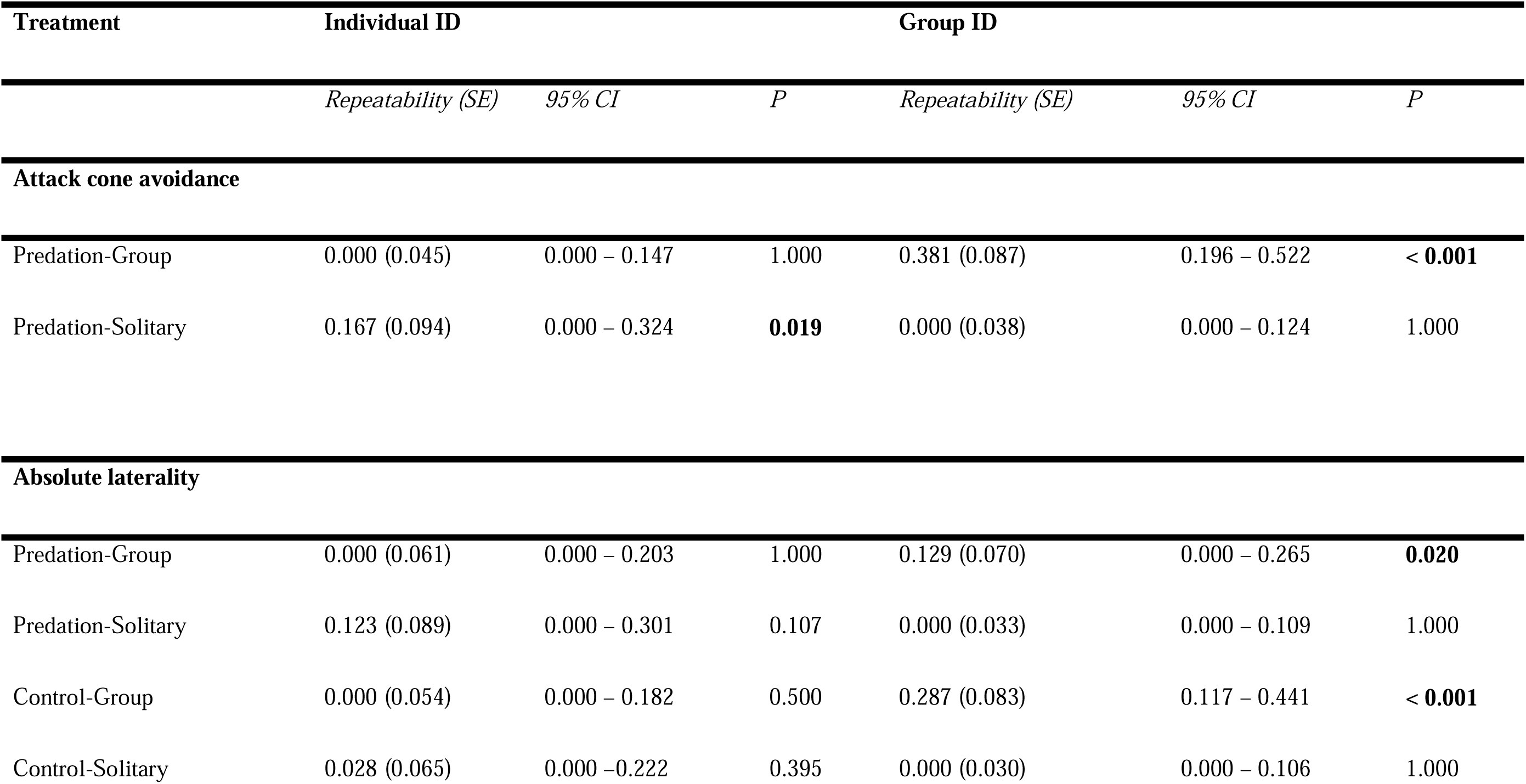
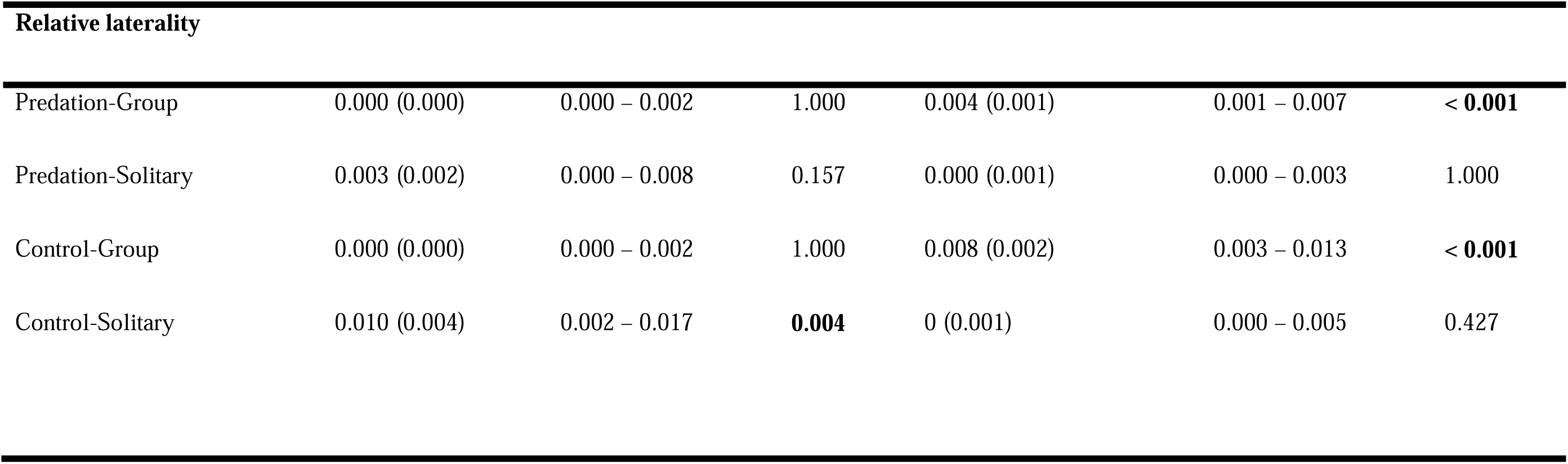
Repeatability of behavioural parameters (attack cone avoidance, absolute laterality, and relative laterality) of female P. reticulata in each of the four treatments: “Predation-Group”, “Predation-Solitary”, “Control-Group”, and “Control-Solitary”. Values in bold are deemed to be statistically significant.

Solitary” treatment. Group ID was not found to be repeatable in the solitary trials nor individual ID in the group trials (*Table 1*).

### Absolute lateralisation

Low levels of absolute lateralisation were found across all treatments. A large proportion of the trials, 98.94%, were below an absolute laterality index of 0.5, with 58.38% below 0.1. Despite the low levels, several factors were found to have a statistically significant impact on absolute lateralisation. Solitary trials produced significantly higher absolute lateralisation indexes (mean absolute laterality index ± SD: 0.13 ± 0.12) compared to group trials (0.09 ± 0.09: *Figure 3*: GLMM: χ^2^_(1)_ = 51.79, p < 0.001). Control trials demonstrated higher absolute lateralisation indexes (0.12 ± 0.11) compared to predation trials (*Figure 3*: 0.10 ± 0.10: χ^2^_(1)_ = 5.06, p = 0.024). Guppy activity (cm/min) was positively correlated with absolute laterality index (χ^2^_(1)_ = 32.74, p < 0.001). Analysing the predation trials with the inclusion of predator-specific parameters in the model did not qualitatively alter these results, with both the grouping treatment (χ^2^ = 28.90, p < 0.001) and guppy activity (χ^2^ = 23.62, p < 0.001) being the only explanatory variables significantly impacting absolute laterality.

**Figure 3:**
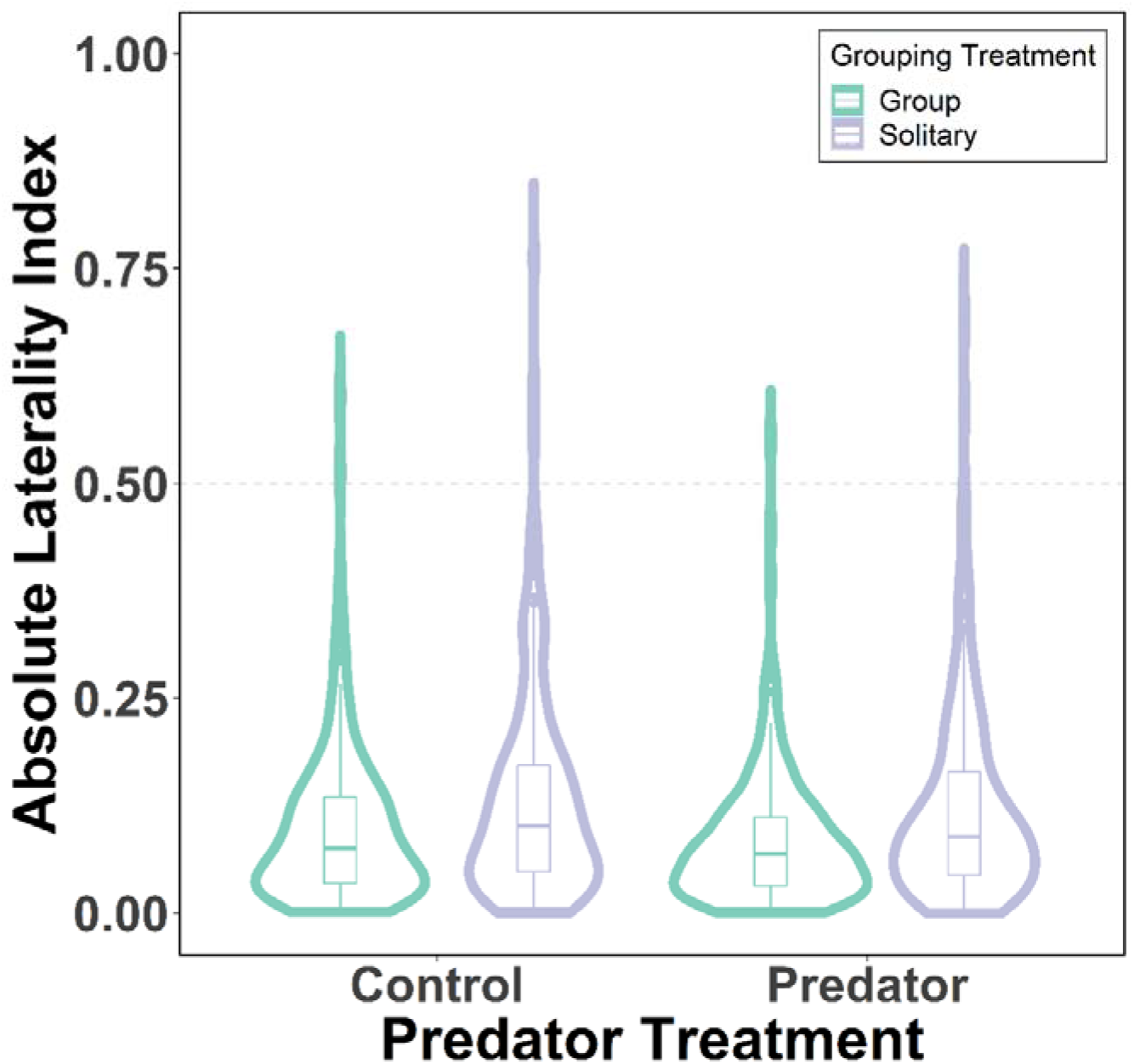
Absolute (intensity) lateralisation index for the predation and grouping treatments. Medians are illustrated by the thick horizontal central line, the boxes enclose the interquartile ranges (IQR), and the whiskers represent cases within 1.5□×□IQR. The circles represent datapoints outside of the whiskers. The data distributions are also represented with kernel density plots.

Repeatable absolute laterality indexes were observed for the group ID random effect when data from all treatments were included in the analysis (LMM: R = 0.050, 95% CI = 0 – 0.112, N_obs_ = 752, N_grp_ = 44, p = 0.012); however, individual ID was only marginally repeatable (R = 0.039, 95% CI = 0 – 0.099, N_obs_ = 752, N_ind_ = 99, p = 0.052). When assessing treatments separately, group ID was significantly repeatable within both predation and control group trials (*Table 1*) but not within the solitary trials. Absolute lateralisation for individual ID was not repeatable in any of the treatments (*Table 1*: p = 0.107 – 1).

### Relative lateralisation

Low levels of relative lateralisation were identified in all treatments. However, there was a significant leftward bias in viewing the central chamber when the predator was present (mean relative lateralisation index: -0.01 ± 0.14) compared to the control trials (*Figure 4*: 0.01 ± 0.16, GLMM: χ^2^_(1)_ = 4.22, p = 0.040). Trials later in the series also had a significant leftward trend (χ^2^_(1)_ = 4.01, p = 0.045). Relative lateralisation was not different between the solitary and group treatments (*Figure 4*: χ^2^_(1)_ = 0.09, p = 0.760). Analysis of the predation treatment including the additional predator-specific parameters bore no significant predictors of relative lateralisation (Grouping treatment: χ^2^_(1)_ = 2.25, p = 0.133).

**Figure 4:**
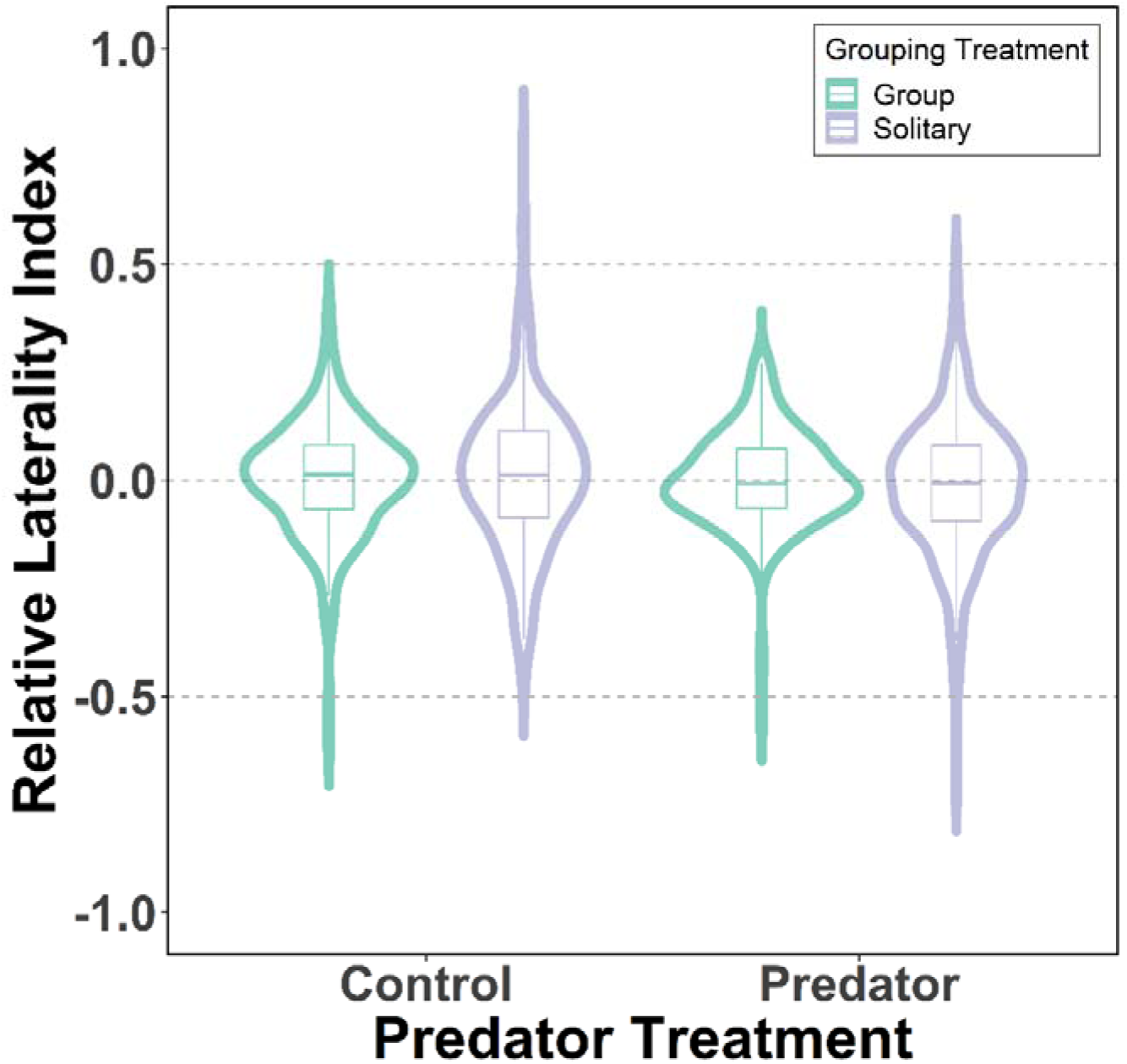
Relative (directional) lateralisation index for the predation and grouping treatments. Medians are illustrated by the thick horizontal central line, the boxes enclose the interquartile ranges (IQR), and the whiskers represent cases within 1.5L×LIQR. The data distributions are also represented with kernel density plots.

An individual’s relative laterality was repeatable when including all treatments (GLMM: R = 0.003, 95% CI = 0.001 – 0.005, N_obs_ = 752, N_ind_ = 99, p < 0.001), as well as within each group (R = 0.002, 95% CI = 0 – 0.003, N_obs_ = 752, N_grp_ = 44, p = 0.019). Investigating repeatability of relative lateralisation within each treatment separately identified, again, that group ID was significantly repeatable within the predation and control group treatments (*Table 1*), and this was not maintained in the solitary treatments (*Table 1*). Individual ID was significantly repeatable within the “Control-Solitary” treatment (*Table 1*) but was not repeatable within any of the other treatments (*Table 1*).

## Discussion

Previous studies have found that visual lateralisation is enhanced in response to a predatory threat (Bisazza et al., 1997a; Bisazza, Facchin, et al., 1998)L or when shoaling (Brown et al., 2007; Dadda et al., 2012; Sovrano et al., 1999)L. There are benefits expected in being able to observe both stimuli simultaneously (Dadda & Bisazza, 2006a; Rogers et al., 2013)L. Being able to monitor conspecifics while simultaneously observing predators would minimise detection latency and neural processing time, maximising response efficiency and group synchronisation (Bisazza et al., 2000; Brown et al., 2004; Vallortigara, Regolin, et al., 1999)L. It is evident in our study that the guppies perceived the blue acara as a predatory threat, with a clear avoidance of the predator’s attack cone (Magurran & Seghers, 1990)L. Despite this, absolute visual lateralisation was not enhanced when viewing the predator nor when shoaling with a group. Notably, absolute lateralisation was also not enhanced when simultaneously performing these activities as there was no significant interaction between these treatments. However, we did find a statistically significant, albeit small, preference for left eye use when facing a predator in the central chamber across both group and solitary trials.

In our study, we observed low levels of absolute lateralisation across all experimental conditions. Control trials, however, showed higher absolute laterality than trials containing the live predator, contrary to predictions based on previous studies (Brown et al., 2004; Broder & Angeloni, 2014; Facchin et al., 1999; Sovrano et al., 1999)L. Whilst there are potential benefits of visual lateralisation, its presence under predation threat or in high predation environments has not been consistently found. Strong visual lateralisation might be disadvantageous when facing predators because of the high number and diversity of predators, making prey species vulnerable to attacks from multiple directions (Rogers, 2000; Rogers et al., 2004)L. The population used within this experiment were derived from a high predation environment but had no prior experience with predators for multiple generations. Nonetheless, these fish demonstrated clear attack cone avoidance (Magurran & Seghers, 1990)L. The reduced lateralisation indexes demonstrated in the predation trials could, in fact, be their response to the movement of the predator to avoid the predator’s attack cone, rather than their movement being primarily to determine with which eye to observe the predator.

Despite the lower absolute lateralisation found in our predator trials, guppies did demonstrate a significant leftward bias compared to control trials. Information from the left eye is primarily processed by the contralateral right hemisphere of the brain. In fish, the right hemisphere is primarily associated with vigilance, fear responses, and predator detection (Cantalupo et al., 1995; Facchin et al., 1999; Sovrano et al., 2001; Vallortigara and Rogers, 2005), while the left hemisphere is linked to routine behaviours like feeding and social interactions (Bisazza et al., 1997b; Miklosi et al., 1999; Rogers et al., 2013). Previous research has similarly identified a trend in right eye-use (left turning) in goldbelly topminnows (*G. falcatus*) (Facchin et al., 1999)L and male eastern mosquitofish (*G. holbrooki*) when observing a simulated/dummy predator (Bisazza et al., 1997a)L. It may be that even a slight directional bias, as found in our study, may give lateralised individuals an advantage when under threat of predation. Even small differences between individual responses can be the difference between life and death when under predation threat (Kelley & Magurran, 2003; Lima & Dill, 1990)L. While our study identified significant differences in visual lateralisation between treatments, these differences are modest compared to other lateralisation studies (e.g. Brown et al., 2004; Facchin et al., 1999; Hulthén et al., 2021). The low levels of lateralisation observed across all treatments, despite some statistically significant effects, raise questions about their biological relevance. Therefore, the implications in real-world scenarios may be limited and these findings suggest the need for further investigation to uncover whether there is indeed biological significance and potential adaptive advantages.

Absolute lateralisation was found to be significantly higher when fish were tested alone compared to when they were tested in groups, contrary to our prediction. As a shoaling species, solitary individuals may experience a higher level of perceived threat, stress or fear when alone, regardless of there being a predator present. Stress has been found to be a key factor in determining the degree of behavioural lateralisation, with higher levels of stress contributing to higher levels of behavioural lateralisation (Berlinghieri et al., 2021; Halpern, 2005; Ocklenburg et al., 2016)L. It is possible that solitary individuals would make better use of visual lateralisation to monitor their surrounding environment for predators and completing other tasks, such as foraging, while groups may gain sufficient protection from collective vigilance (Ward et al., 2011)LL. However, despite the importance of the social context for lateralisation, most studies have only assessed solitary individuals (Brown et al., 2004)L.

In our study, a significant interaction between the grouping treatment and predator presence was not identified. However, previous work has identified relationships in visual lateralisation for anti-predatory and social contexts. Despite no population-level asymmetric eye-use, in two separate assays goldbelly topminnows (*G. falcatus*) preferred to view a predator stimulus and their ‘social’ reflection with different eyes (Dadda et al., 2012)L. Comparably, *Xenophallus umbratilis* demonstrate a clear eye side-bias when observing either a predatory or social stimuli, with left morphs consistently using their right eye for both, and right morphs using their left (Johnson et al., 2020). Female *B. episcopi* from high predation environments have shown increased levels of absolute lateralisation, and a trend in right eye-use (left turn), when viewing a conspecific in a detour test compared to those from low predation environments. This trend was also maintained and enhanced in laboratory-reared offspring (Brown et al., 2007)L. Female *B. episcopi* from high predation areas assessed in groups demonstrated a right eye-use preference when viewing a live potential predator (*A. pulcher*), while low predation individuals had no preference (Brown et al., 2004)L.

Previous assessments have found mixed results regarding the repeatability of lateralisation in solitary individuals (Penry-Williams et al., 2022; Roche et al., 2020; Vinogradov et al., 2021)L. Roche et al. (2020) found relative lateralisation in five fish species was not repeatable in a detour assay, including in feral guppies (*P. reticulata*) (Irving & Brown, 2013)L, despite many individuals demonstrating a strong directional bias within a trial. Similarly, limited consistency in relative and absolute lateralisation was found between individuals’ trials in three-spined sticklebacks (*Gasterosteus aculeatus*) (Panizzon et al., 2024)L. Repeatability in relative lateralisation was found to be higher in guppies across various contexts, but absolute lateralisation was only repeatable in males (McLean & Morrell, 2020)L. Similarly, Vinogradov et al. (2021)L found that relative laterality in female eastern mosquitofish (*G. holbrooki*) was significantly repeatable in five of six treatments incorporating social and control stimuli. Repeatability of lateralisation in our study was observed in the group-level random effect across both predation and control treatments. However, there was limited evidence for repeatability at the level of the individuals. This result suggests that social processes, primarily social conformity, may have a greater effect than predation risk on determining the variation observed in lateralisation (Brown & Laland, 2002; Ioannou & Laskowski 2023)L.

A number of studies have demonstrated among-group behavioural variation in fish shoals, even when groups are formed of randomly-selected individuals (Jolles et al. 2018) or group membership is designed to minimise inter-group variation (MacGregor & Ioannou 2021). The manipulation of shoal compositions based on body size to allow for individual identification in our study may have had an impact on the results. Forming shoals of individuals that are more diverse in body size should have increased within-group variation in behaviour relative to among-group variation. This should have reduced the differences among groups, and hence reduce the repeatability in the group identity random effect (Ioannou & Laskowski, 2023)L. Thus, the strength of social conformity in lateralisation may be underestimated, and social conformity may be playing a major role in determining eye use in social species. Consistent with this, in a study of sixteen fish species, shoaling species (e.g. Poeciliidae and Cyprinidae) were more likely to demonstrate a population-level conformity in the direction of lateralisation when undertaking predator escape responses, while non-shoaling species (e.g. freshwater Gobiidae and *Ancistrus* sp.) were more likely to demonstrate a mixture of individuals with both right and left alignment (Bisazza et al., 2000)L.

In our study, it was hypothesised that if lateralisation is an important anti-predation mechanism, then the presence of a predator should reduce the variability between individuals in this trait in order to maximise survival under perceived predation threat (Toscano et al., 2014). In agreement with this hypothesis, repeatability in relative lateralisation was identified in the solitary trials at the level of the individual in the control trials, but not in the trials with a predator stimulus. Alternative anti-predation mechanisms may also be adopted by solitary individuals in the predator trials, such as “protean” movements, which may lead to unpredictable swimming behaviour and, therefore, measured eye-use (Jones et al., 2011; Szopa-Comley & Ioannou, 2022)L. Not mutually exclusively, the movement of the guppies in the predator trials could have been responding more to avoiding the attack cone of the predator, rather than their movement being primarily to determine with which eye to observe the predator. Behaviours more sensitive to environmental or motivational influences may be less predictable and repeatable (Bell et al., 2009)L, such as those influenced by energetic needs (MacGregor et al., 2021)L, social interactions (MacGregor & Ioannou, 2021; Rands & Ioannou, 2023),L or ecological variables (Castellano et al., 2002; Roy & Bhat, 2018; Smith & Hunter, 2005)L.

In conclusion, this investigation provides evidence suggesting that visual lateralisation is not enhanced under predation threat or in social groups, contrary to our predictions, with relatively low lateralisation indexes observed across all treatments. Repeatability of lateralisation indexes with the group-level random effect was identified but limited repeatability at the individual level suggests an important role of social conformity in lateralisation. We suggest that these results indicate social processes may have a greater effect than predation risk on variation in lateralisation. Given the predictions of lateralisation generally suggest greater benefits to groups over solitary individuals, we suggest that a greater consideration of groups when assessing lateralisation in addition to individuals is required to disentangle the potential benefits of visual lateralisation.

## Acknowledgements

This research was supported through the NERC (Natural Environment Research Council) GW4+ Doctoral Training Partnership and Macquarie University Cotutelle (NE/L002434/1). We would like to thank Tomos Potter, Anja Felmy, and the Guppy Project at the University of Oxford for the collection, breeding, and transport of guppies used within this research. We would also like to thank Robert Heathcote, Jolyon Troscianko, and collaborators for access to their custom ImageJ plugin for assessing fish orientation, originally presented in Heathcote et al. (2020). We also thank Anne-Kristin Lenz for helpful discussion on extraction protocols and calculations.

## Author Contributions

Iestyn L. Penry-Williams: Conceptualization (lead), Data Curation (lead), Formal Analysis (lead), Investigation (lead), Methodology (lead), Project Administration (lead), Visualization (lead), Writing – Original Draft Preparation (lead).

Culum Brown: Funding Acquisition (supporting), Supervision (supporting), Writing – Review & Editing (lead).

Christos C. Ioannou: Conceptualization (supporting), Funding Acquisition (lead), Methodology (supporting), Project Administration (supporting), Resources (lead), Supervision (lead), Writing – Review & Editing (supporting).

## Data Availability Statement

The data for the analyses are provided as Supporting Information.

